# Exosomal miRNA-124 is involved in the immune regulation of cervical cancer by regulating CD45 alternative splicing

**DOI:** 10.1101/2024.05.06.592634

**Authors:** Minxing Liang, Man Yi, Yuyang Yuan, Chengyi Chen, Qiyu Wang, Bo Lan, Jun Yang, Xin Li, Yue Yin

**Affiliations:** School of Pharmacy, Guilin Medical University; Department of Gastrointestinal Surgery, the Affiliated Hospital of Guilin Medical University; School of Basic Medicine, Guilin Medical University; Department of the Functional Science laboratory, Key Laboratory of Tissue Microenvironment and Tumor, Guilin Medical University; Department of Histology and Embryology, Guilin Medical University, Guangxi, 541199, Guilin, China

**Keywords:** Exosomes, mRNA, Cervical cancer, Regulation of immunity

## Abstract

**Objective:** To investigate the regulatory mechanism of miRNA-124 in tumor immune regulation of cervical cancer.

**Methods:** Peripheral blood samples of cervical cancer patients and transient infection controls were collected to extract exosomal miR-124 and Stem-loop Q-PCR to detect the expression level of miR-124. QPCR, WB, and flow cytometry were used to detect the mRNA and protein expression of Th cell differentiation and function-related genes. Transfection, QPCR, and flow cytometry were used to detect the effects of miR-124 overexpression and silencing on the differentiation of Th immune memory cells. In vitro, cell-killing experiments were performed to detect the tumor-killing activity of TH immune memory cells co-cultured with the Hela cell line, and the downstream target genes of miR-124 were predicted and verified by luciferase. Multiplex PCR was used to detect the genotype of miR-124 SNP in peripheral blood DNA. QPCR and WB were used to detect the mRNA and protein expression and phosphorylation of miR-124 target gene GSK3-PSF-CD45 pathway, and GSK3 inhibitor was used to block the pathway.

**Results:** We found that exosomal miRNA-124 was significantly downregulated in cervical cancer tissues and affected Th cell immune function and memory cell generation. The overexpression of miRNA-124 can directly target GSK3, phosphorylate the target gene, and inhibit the expression of GSK3, thereby increasing the expression of CD45, increasing the ability of Th immune cells to recognize and kill the HPV virus, and significantly inhibiting the proliferation and migration of cervical cancer cells.

## Introduction

China has about one-third of the world’s cervical cancer burden, with about 98,900 new cases of cervical cancer in 2015, and the incidence is expected to increase year by year^[1]^. Human papillomavirus (HPV) is the main pathogenic microbe that causes cervical cancer. Studies have found that HPV DNA can be detected in 99.7% of cervical cancer tissues^[2]^. So far, HPV has been found to exist in more than 200 subtypes, according to the epidemiological correlation with cervical cancer divided into high-risk type and low-risk type; Among them, the most common high-risk HPV types, 16 and 18, cause about 70% of cervical cancer^[2]^. The common clinical manifestation is cervical intraepithelial neoplasia (CIN), and CIN grade progresses from CIN1 to CIN2 and CIN3 after several years. Without regular screening and timely treatment, high-grade CIN may develop into cervical cancer years or decades later^[2, 3]^. Although the prevalence of HPV infection is high in the population, the majority of infections are effectively cleared by the immune system. Only a small proportion of individuals with HPV infection develop persistent infections that eventually lead to cervical cancer, and the underlying contributing factors remain incompletely understood. Research has demonstrated significant associations between genetic polymorphisms in immune function-related genes (such as MAGI-3, SMAD3, and HLA-G) and HPV clearance among patients co-infected with HIV, suggesting that host regulation of immune response may play a role in the body’s ability to respond to and clear HPV^[4]^. Additionally, studies have indicated that genetic susceptibility to cervical cancer is linked to genes encoding cytokines or receptors closely associated with the Th-cell immune response (CD95, IL-10, IL-4, IL-6, TNFA, IFNG)^[5, 6]^. Therefore, investigating the interaction between genes involved in regulating immune response during the occurrence and progression of cervical cancer holds great significance and practical value for exploring potential molecular targets for future disease prevention and treatment strategies as well as prognostic monitoring methods.

Exosomes are a type of membranous vesicles, ranging from 20 to 100nm in diameter, that are released by cells into the extracellular matrix. They carry proteins, DNA, and non-coding RNA, and participate in various physiological processes such as immune response, antigen presentation, and intercellular communication^[7]^. By transporting antigens to dendritic cells, exosomes can induce the immune response of CD8+ T cells^[8]^, express immune checkpoint ligand PD-L1 and apoptosis ligand FasL to induce lymphocyte apoptosis^[9-11]^, and promote T cell function to inhibit NK cell and T cell activity. Previous studies have demonstrated that exosomes derived from HPV-positive cell lines can enhance dendritic cell maturation, suggesting their positive role in anti-tumor immunity^[12]^. However, exosomes derived from nasopharyngeal carcinoma cells can induce apoptosis of specific CD4+ T cells infected with EBV (Epstein-Barr virus) in nasopharyngeal carcinoma patients and suppress the cellular immune effect of Th1 (Type 1 helper T cells)^[13]^. Additionally, exosomes can exert their effects through the miRNAs they carry. Studies have revealed that exosomes released by Hela cell lines derived from cervical cancer contain various miRNAs. The expression profiles of these miRNAs change when oncogenes E6/E7 are silenced, indicating the potential involvement of exosomes in tumor proliferation regulation^[14, 15]^.

The microRNA (miRNA) is a small RNA molecule derived from the primary miRNA (pri-miRNA), which undergoes further processing to form pre-miRNA with a length of 300-1000 nt, and eventually generates small RNA molecules of 20-22 nt. These small RNAs bind complementarily to the 3’ UTR region sequence of target gene mRNA, leading to subsequent degradation or translation inhibition^[11, 16]^. As mentioned earlier, exosomes also contain miRNAs and play a crucial role in viral immunity. Recent studies have identified miR-124 as an important regulator of immune response and inflammatory response, suggesting its potential as a diagnostic marker and therapeutic target for future treatments^[17, 18]^. In HPV-related cohort studies, it was observed that HPV patients over the age of 30 who tested negative for methylation in both miR-124 and FAM19A4 genes had significantly reduced risk of cervical cancer. Preliminary clinical diagnostic tests have indicated that these markers could be used for screening high-grade CIN^[19, 20]^. Furthermore, research has shown that miR-124 is associated with the development of cervical cancer caused by HPV infection and the prognosis of HPV-induced precancerous lesions^[21, 22]^. Therefore, further investigation is needed to understand the relationship between miR-124 and persistent HPV infection leading to cervical cancer along with its underlying mechanism.

## 1 Materials and Methods

### 1.1 Materials

#### 1.1.1 Experimental materials

Peripheral blood samples of cervical cancer patients and transient infection control population were collected, which were required to be at least 1.5ml in volume and stored in an EDTA anticoagulant tube, and then divided into EP tubes at minus 80□after full anticoagulation. Culture of Hela cell line for cervical cancer, etc.

#### 1.1.2 Experimental apparatus

Fluorescence quantitative PCR instrument, low temperature and high-speed centrifuge, electrophoresis instrument, ultrasonic fragmentation instrument, electro revolution instrument, automatic gel imaging analysis system, flow cytometer, CO2 incubator, inverted microscope, frozen microtome, paraffin microtome, fluorescence microscope, image acquisition and analysis system, and other cell biology research equipment, etc.

#### 1.1.3 Experiment reagents

Qiagen exosome extraction kit, RNA extraction kit, Trizol, etc.

### 1.2 Methods

#### 1.2.1 Extraction of exosome miR-124 and Stem-loop Q-PCR to detect the expression level of miR-124

Peripheral blood samples were collected from cervical cancer patients and transient infection controls, and the plasma was separated by centrifugation and frozen at -80 □. Plasma exosomes were separated by the steps of pre-filtration, mixing, column, washing, and elution according to the Qiagen exosome extraction kit. Exosomes were extracted by RNA extraction kit with column extraction method, and miR-124 specific stem-loop reverse transcription primers were synthesized to synthesize cDNA. The expression level of miR-124 in exosomes was detected by Stem-loop Q-PCR.

#### 1.2.2 Cell culture

Healthy human PBMC were isolated by Ficoll density gradient centrifugation and then cultured and activated with anti-CD3 5μg/ml. The exosomes were extracted and eluted in 10ml plasma per 5×105 cells for 48 hours.

#### 1.2.3 QPCR, WB, and flow cytometry were used to detect Th cell differentiation and function-related gene mRNA and protein expression

RNA was extracted using Trizol. The mRNA expression of memory cell differentiation-related genes CD127, CTLA4, KLRG1, and CD44, as well as the mRNA expression of IFNγ, TNFβ, IL-4, and IL-10 in PBMC cells treated with exosomes from patients and controls, was detected by Q-PCR. The expression changes of CD127, CTLA4, KLRG1, and CD44 in cell subsets were analyzed by flow cytometry and Flowjo software. The percentages of CD3+CD4+CD45RA- and CD3+CD8+CD45RA-differentiated memory cells and the expression of these key cytokines were detected by flow cytometry.

#### 1.2.4 The effects of overexpression and silencing of miR-124 on Th immune memory cell differentiation were detected by transfection, QPCR, and flow cytometry

The expressions of CD127, CTLA4, KLRG1 and CD44 at mRNA and protein levels were detected by Q-PCR and flow cytometry. The mRNA expressions of IFNγ, TNFβ, IL-4, and IL-10 in the cells were observed. The percentages of CD3+CD4+CD45RA- and CD3+CD8+CD45RA-differentiated memory cells were detected by flow cytometry. The expression of these key cytokines was observed by flow cytometry membrane breaking staining.

#### 1.2.5 In vitro and in vivo cell killing experiments

PBMC were transfected with miR-124 mimic and inhibitor and co-cultured with Hela cell line to detect the tumor-killing activity in vitro. After co-culture, the tumor cells of each experimental group and the control group were injected subcutaneously into BALB/C nude mice to form tumors. The animal status and tumor size were observed regularly. The animals were sacrificed to take out the tumor weight and measure the size.

#### 1.2.6 min walk website predicted downstream target genes of miR-124 and validated luciferase

The miRWalk website was used to search the target gene list of miR-124 and screen genes related to immune function. Previous experiments had focused on the regulatory protein GSK3 related to CD45 alternative splicing. The online tool RNAhybrid was used to perform bioinformatics analysis of the 3’utr complementarity of miR-124 and GSK3.

The mRNA and protein expression levels of GSK3 were detected by Q-PCR and Western blot after RNA extraction from peripheral blood mononuclear cells. The expressions of GSK3 mRNA and protein in PBMC were detected by Q-PCR and Western blot after transfected with miR-124 mimic and inhibitor. Luciferase reporter system was used to construct gsk3-3’utr reporter plasmid and co-transfection with miR-124 mimic was used to detect the targeting regulation of miR-124 on GSK3. A complementary double mutation experiment was used to verify the specific complementary binding effect of miR-124 and GSK3.

To confirm the relationship between miR-124 and 3’utr target gene GSK3, Normal and mutant Luciferase reporter gene cell models (pMIR-REPORT luciferin-gsk3-3’utr and pMIR-REPORT luciferin-mut-gsk3-3’utr) need to be constructed. The complementary sequence of miR-124 in gsk3-3’utr was synthesized and inserted into the 3’utr end of pMIR-REPORT Luciferase by DNA ligase. Hela cell lines were transfected with miR-124 mimic or antagomir, Luciferase reporter plasmid, and Rellina transfection efficiency control plasmid, and the cells were collected to detect the fluorescence value using the luciferase reporter system.

#### 1.2.7 Multiple PCR was used to type miR-124 SNP in peripheral blood DNA

SNPs targeting miR-124 gene: The oligonucleotide of rs531564 (C/G), rs73662598 (A/G/T), and rs73246163 (C/T) of pre-miRNA124 was synthesized and the corresponding restriction sites were designed, annealed and connected with the plKO-mir124 vector (the pLKO plasmid was generously donated by Professor Zhang Hui, Institute of Viruses, Sun Yat-sen University). After screening, the successfully connected expression vectors were obtained. The expression vectors of different genotypes of pLKO-miR124 and lentivirus packaging plasmids psPAX2 and pMD2.G were co-transfected into HEK-293T cells to package the virus, and the virus solution was collected for 4-5 days. Peripheral blood samples were collected in EDTA anticoagulated tubes. PBMCS were separated by Ficoll human lymphocyte separation solution, cultured, and activated with anti-CD3 5μg/ml. Exosomes in the cell culture medium were isolated from infected PBMC according to the Qiagen exosome isolation kit. Total RNA was extracted from cells by Qiagen RNA extraction kit, and the relationship between miR-124 expression level and genotype was detected by Stem-loop PCR and Q-PCR. And the expression levels of the original product, precursor, and mature version of miR-124 in exosomes and in cells, respectively.

#### 1.2.8 mRNA and protein expression and phosphorylation of miR-124 target gene GSK3-PF-CD45 pathway were detected by QPCR and WB

Human PBMC were isolated by Ficoll and transfected with miR-124 mimic and inhibitor by Lipo reagent and cultured for 48 hours. According to the variable shear area of CD45 exon 4 5 ‘end and exon 6 3’ end sequences design across exons primer 5 ‘-GGCAAAGCCCAACACCTTCCCCCACTG - 3’, the downstream use public primers 5 ‘-CTGAAACTTTTCAACCCCTGGTGGCAC - 3’, Polymerase chain reaction (PCR) was used to detect CD45 alternative splicing products. Polyacrylamide gel electrophoresis was used to analyze the proportion of CD45 alternative splicing forms CD45RA and CD45RO in PBMC infected with different genotypes. The expression levels of PSF and TRAP150 mRNA in PBMC infected with different genotypes were detected by Q-PCR. The activity of PSF protein phosphorylation was determined by Western blot.

#### 1.2.9 Detection of GSK3 inhibitor blocking pathway

The GSK3 pathway was blocked by the GSK3 inhibitor, and PBMC was transfected with Mir-124 mimic and inhibitor as above. Alternatively, blood exosomes from cases and controls were isolated as described previously and co-cultured with healthy control PBMCS for 48h. In the above two treatment experiments, PBMCS was collected, and CD45 alternative splicing was detected by electrophoresis of PCR products. The mRNA and membrane protein expression of CD127, CTLA4, KLRG1 and CD44 were detected by Q-PCR and Western blot. The percentages of CD3+CD4+CD45RA- and CD3+CD8+CD45RA-differentiated memory cells were detected by flow cytometry. The mRNA expressions of IFNγ, TNFβ, IL-4, and IL-10 were detected by Q-PCR and Western blot. The expression of these key cytokine membrane proteins was observed by flow cytometry.

## 2 Results

### 2.1 The expression of miRNA-124 in blood exosomes is higher

The miRNA expression profile of exosomes in blood was retrieved by the EVmiRNA database, and the expression of exosomes-derived mirNA-124 in blood was found to be high (FIG. 1). The analysis of miRNA microarray data set of cervical cancer tissues searched by GEO found that the expression of miR-124 in tumor tissues was lower than that of other miRNAs, and there was no significant difference between tumor and normal tissues (FIG. 2). A further search of miR-124 revealed that its distribution in blood-derived exosomes was higher than that in cells of other organs and tissues (Figure 3). These data suggest that miR-124 is more expressed in exosomes, especially in blood exosomes, than in tumors or other normal tissues.

**Figure 1.**
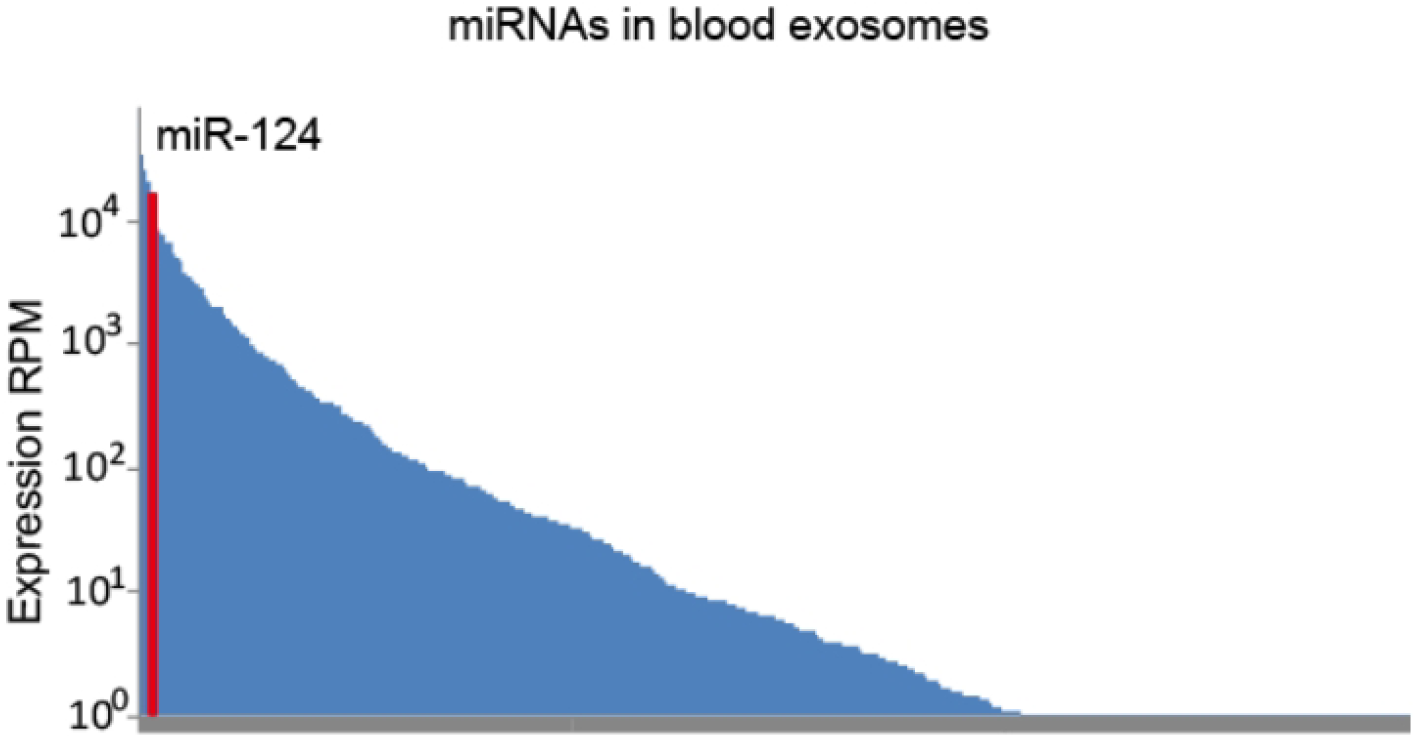
EVmiRNA database (Liu T et al., Nucleic Acids Res, 2019, http://bioinfo.life.hust.edu.cn/EVmiRNA) retrieve blood outside the body secretion expression of miRNAs altogether 586, The red columns are the data of miR-124, RPM, reads per million reads.

**Figure 2.**
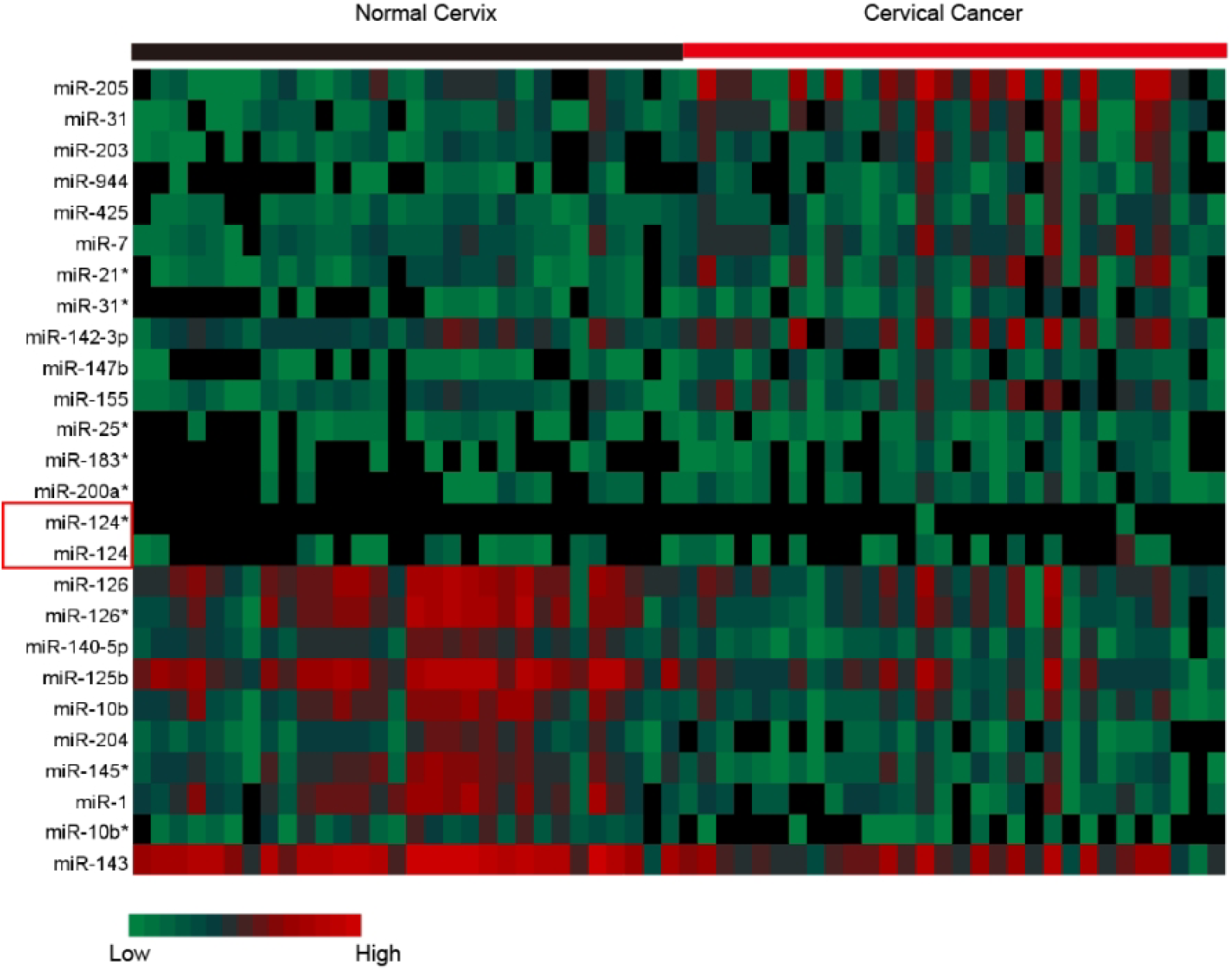
The 24 mirnas in the GSE20592 dataset (Witten D et al, BMC Biol, 2010) that were significantly different between cervical cancer and normal tissues, and the 5p and 3p of the miRNAs of interest included alone.

**Figure 3.**
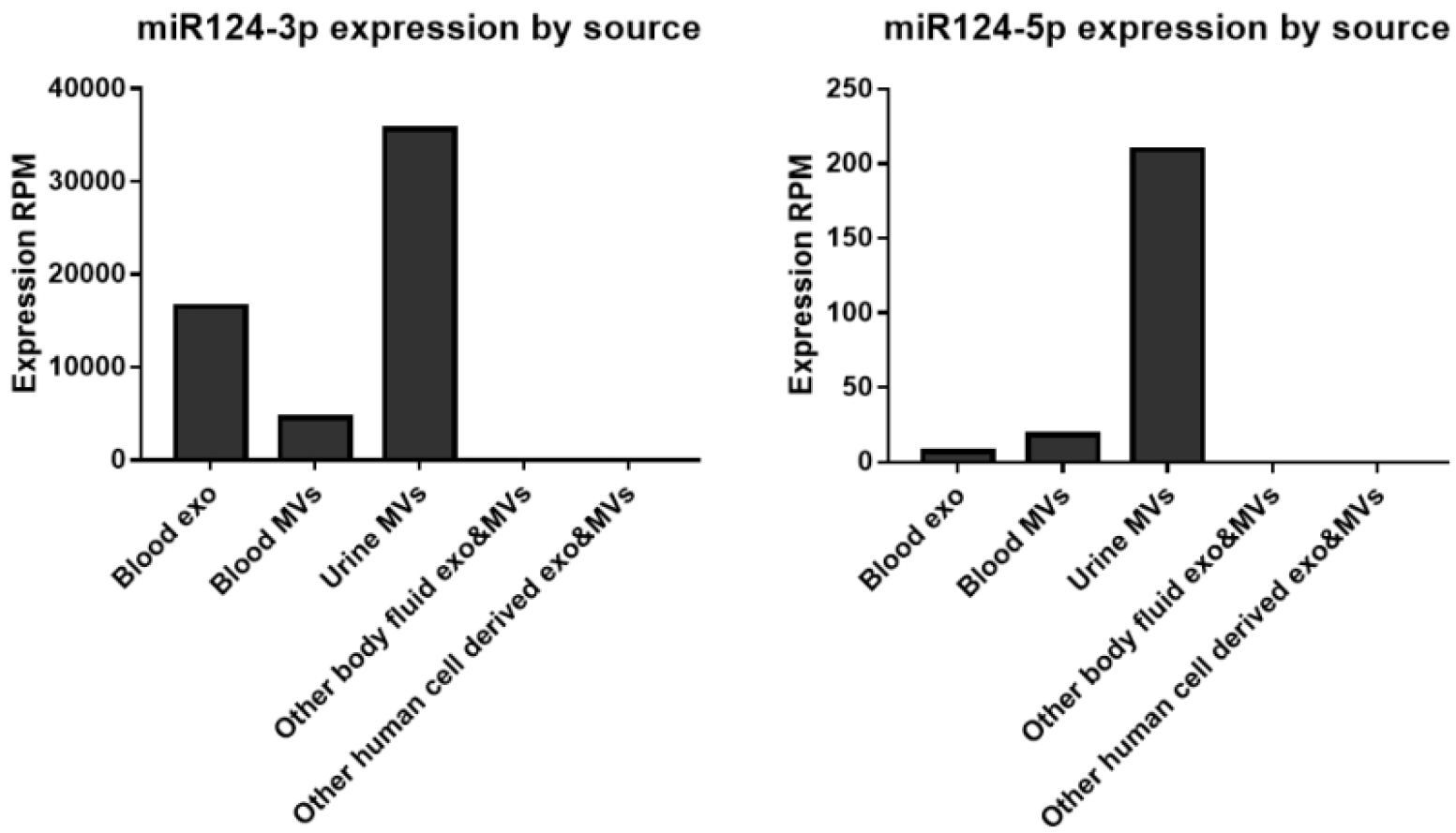
EVmiRNA database retrieval of miR-124 expression in exosomes from multiple tissue sources. A: miR-124-3p; B: miR-124-5p.

### 2.2 Exosomal miR-124 affects the immune function of Th cells and the generation of memory cells

Plasma exosomes extracted from peripheral blood samples of cervical cancer patients and transient infection patients were co-cultured with PBMC of healthy blood donors. It was found that exosomes could reduce the differentiation of CD45RA-memory T cells (Figure 4), and substances in exosomes could regulate the formation of immune memory cells in HPV infection. The stem loop-PCR detection of miR-124 in exosomes found that its expression was significantly reduced in persistent HPV infection (FIG. 5), suggesting that exosomal miR-124 may play a role in regulating Th immune cells in persistent HPV infection. To further find the mechanism of miR-124 regulating Th immunity, we searched GEO datasets through the NCBI public database, and only sample data of cervical cancer tissues and normal tissues were retrieved. Preliminary analysis of microarray data showed that cervical cancer tissues were compared with normal cervical tissues. Fewer than 100 genes were significantly differentially expressed (Figure 6).

**Figure 4.**
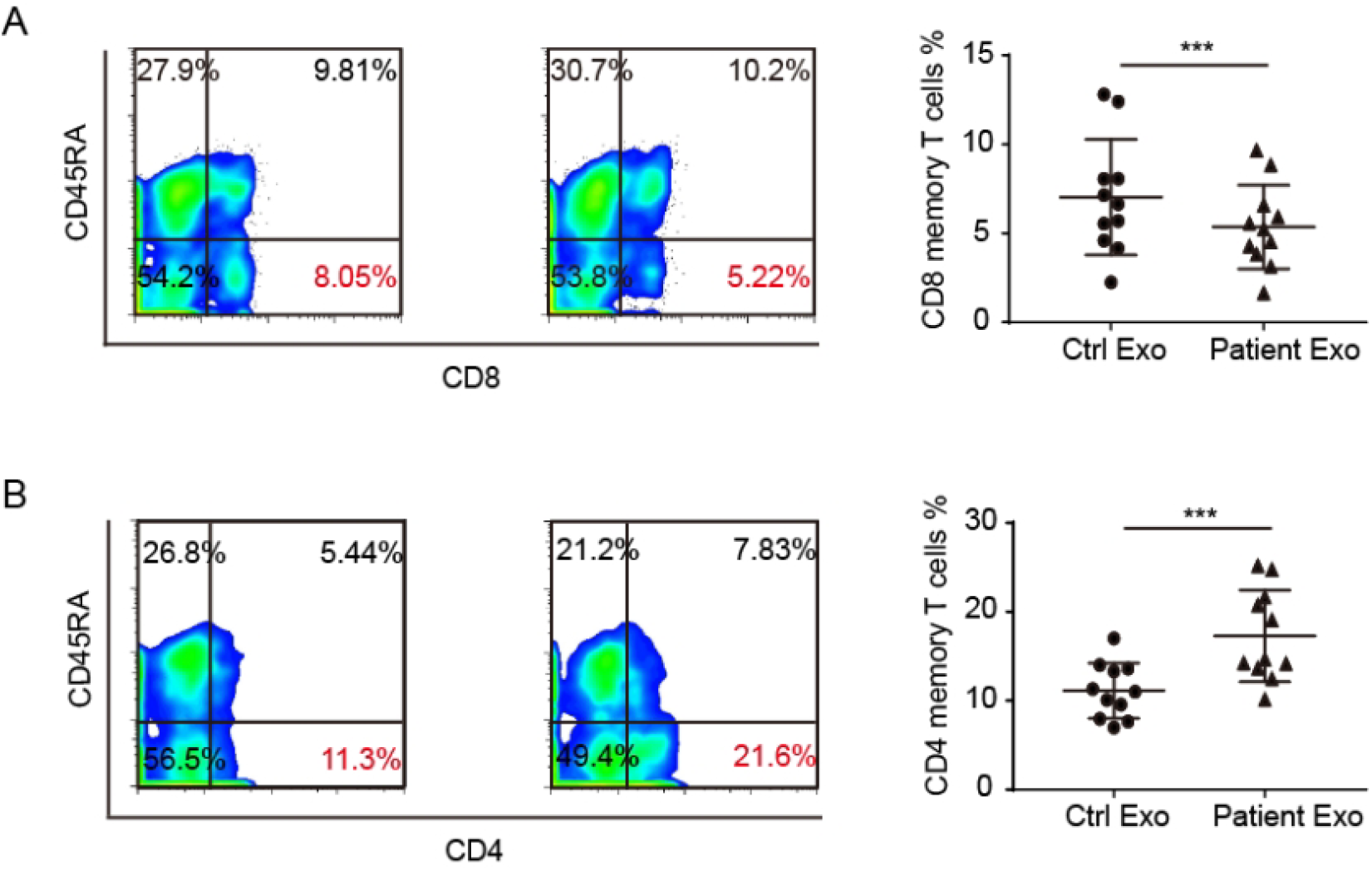
Percentage of PBMC differentiated into memory T cell phenotype detected by exosomes co-cultured with human PBMC for 48 hours, CD45RA-was memory cells, n=3, t-test, ***p<0.001.

**Figure 5.**
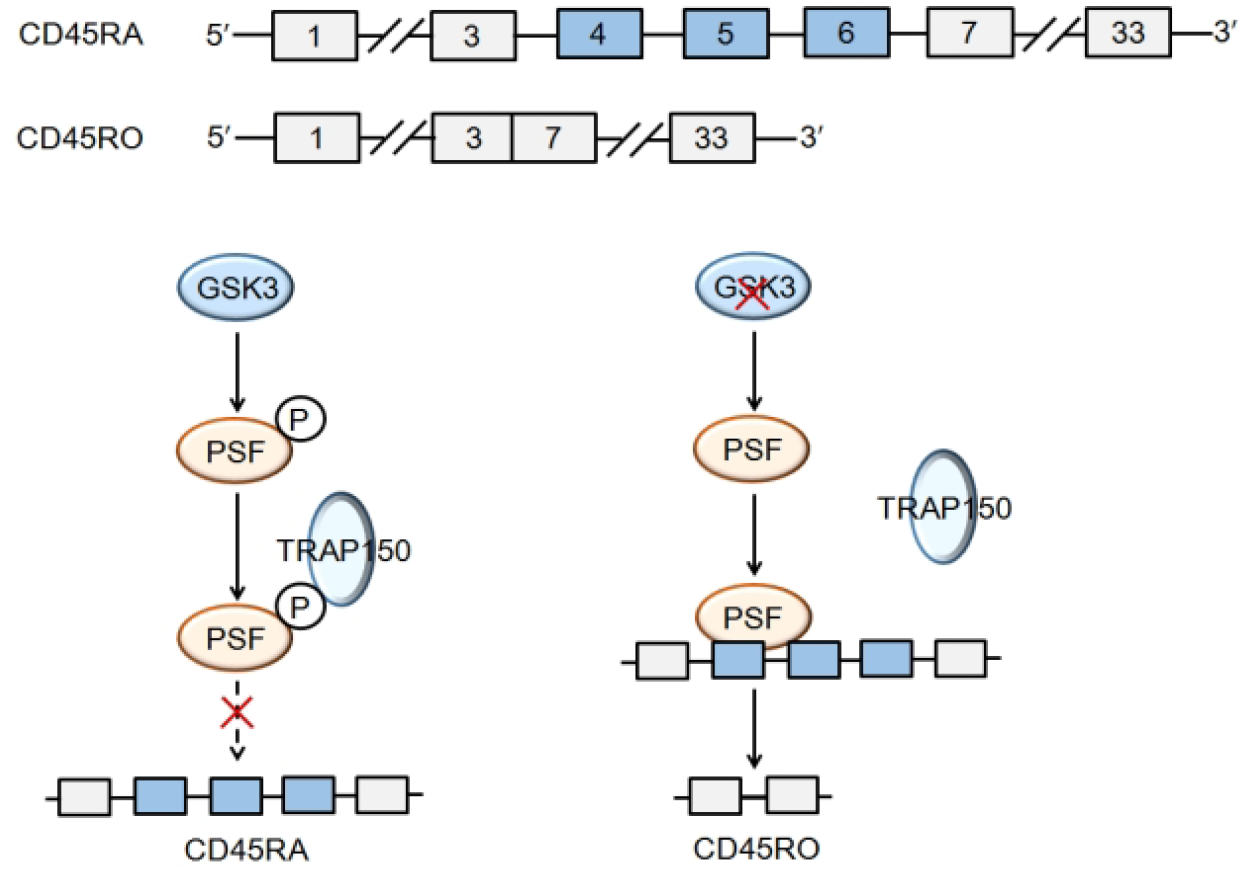
Schematic diagram of CD45RO and CD45RA, two main alternative splicing products of CD45 molecule (top). Schematic representation of the GSK3-PSF-CD45 pathway regulating CD45 alternative splicing (bottom).

**Figure 6.**
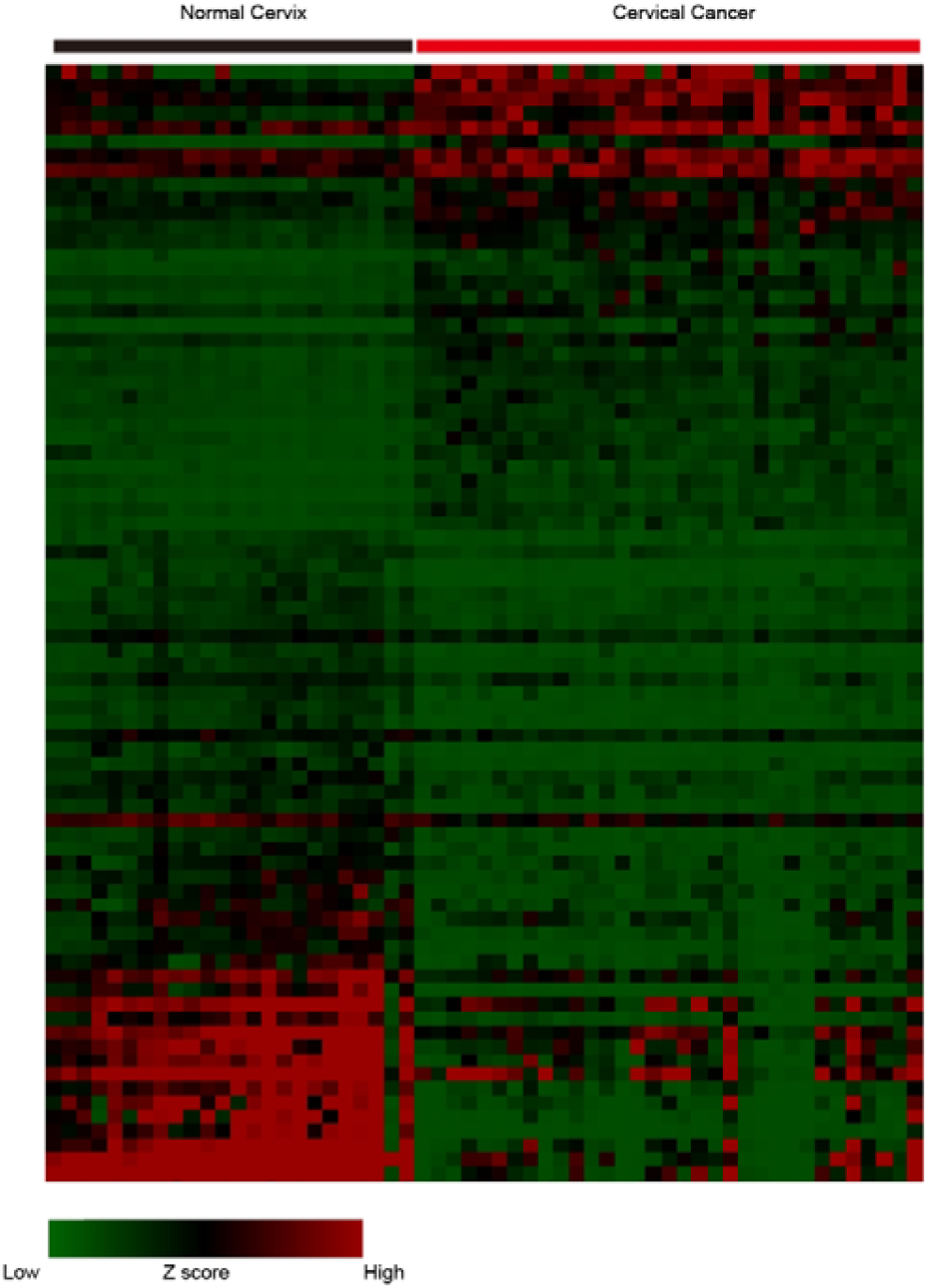
Transcriptome microarray data of 24 non-lesional cervical tissues and 33 cervical Cancer tissues in dataset GSE9750 (Scotto L et al. Genes Chromosomes Cancer 2008). After data bioinformatics analysis, the genes with significant differences were analyzed by heat map (Fold>2, FDR<0.05).

When genes involved in pathways related to Th cell immune responses such as Th1/2 cell differentiation were analyzed separately in all data, it was found that the expression of Th1 cell-related genes was decreased in cervical cancer tissues, while the expression of Th2 cell genes was increased (FIG. 7). Th1 cell immunity plays a role in anti-tumor and antiviral immunity. HPV persistent infection inhibits the immune response by inhibiting the immune function of Th cells to Th1 differentiation. The mRNA extracted from human PBMC was used to detect the expression of Th cell immune-related markers by Q-PCR. It was found that genes related to positive regulation of immune function were down-regulated and genes related to negative regulation of immune function were up-regulated in HPV persistent infection (FIG. 8), which was consistent with the microarray data of pre-cervical cancer. However, the regulatory mechanism of the decreased immune function in HPV infection is still unclear. The gene data related to alternative splicing and memory cell differentiation were also analyzed separately. It was found that the expression of genes related to memory cell differentiation was decreased in cervical cancer tissues, and the expression levels of key genes in the pathway regulating the alternative splicing of memory cell marker CD45, such as GSK3 and PSF, were also altered (FIG. 9), resulting in the reduction of CD45 alternative splicing. Memory cell formation may be reduced. In conclusion, persistent HPV infection may also reduce Th1/2 type differentiation of Th cells and memory cell formation, and this mechanism may be related to the alternative splicing pathway.

**Figure 7.**
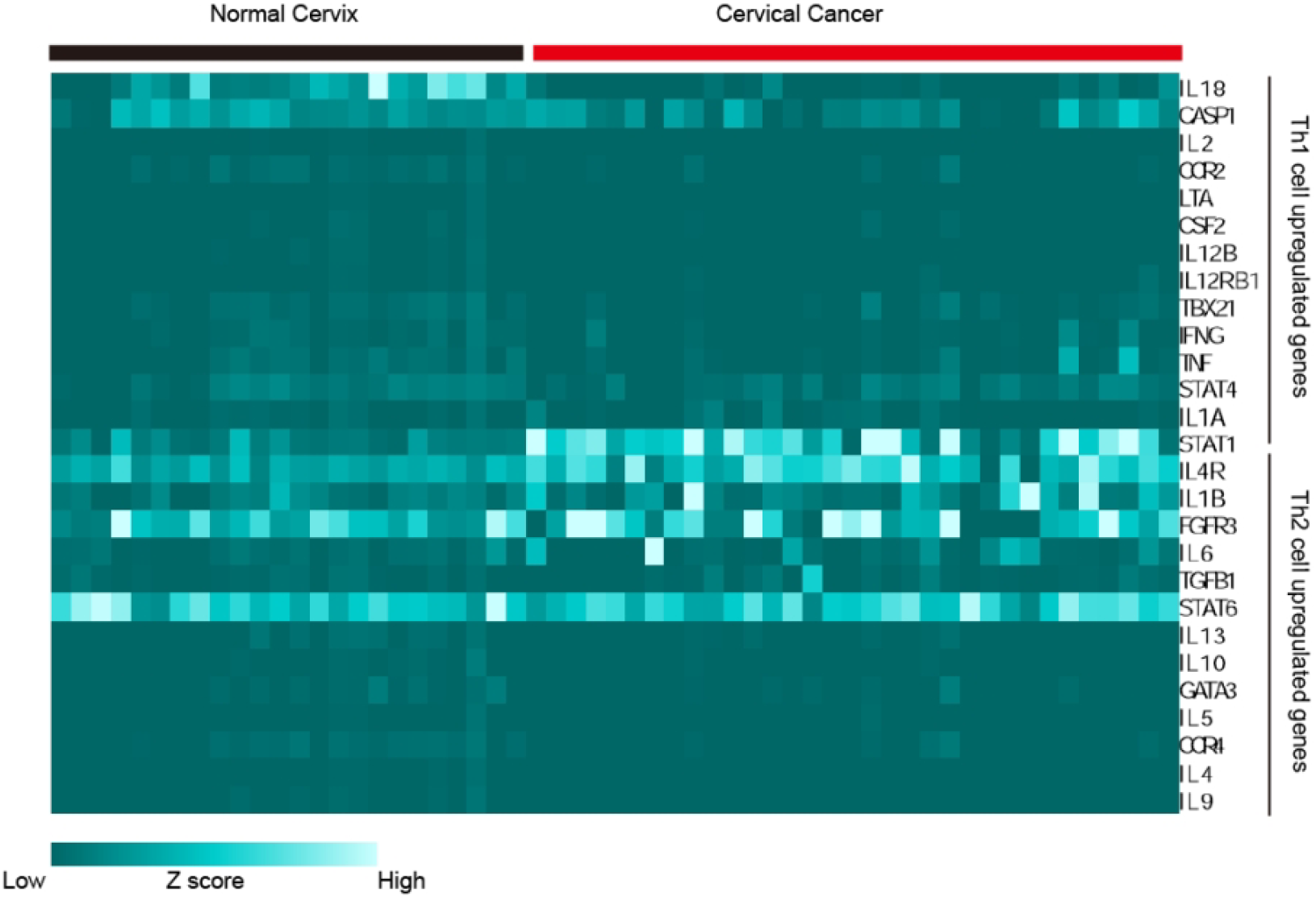
Screening and heat map analysis of marker genes related to Th cell immunophenotyping (Th1, Th2) in the GSE9750 dataset.

**Figure 8.**
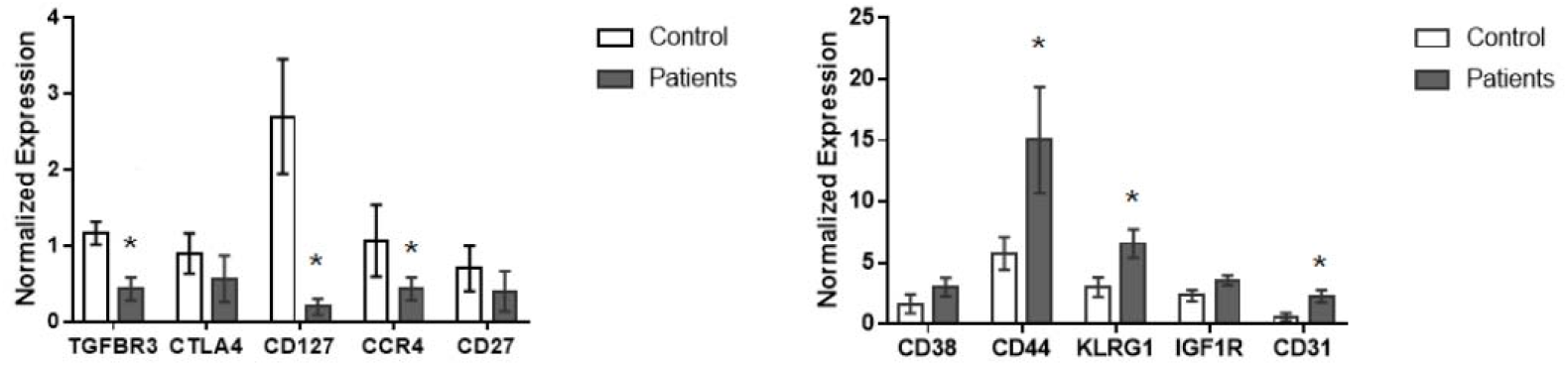
Q-PCR detection of memory T cell-related marker gene mRNA expression in peripheral PBMC cells of cervical cancer patients and transient infection, n=3, t test, *p<0.05.

**Figure 9.**
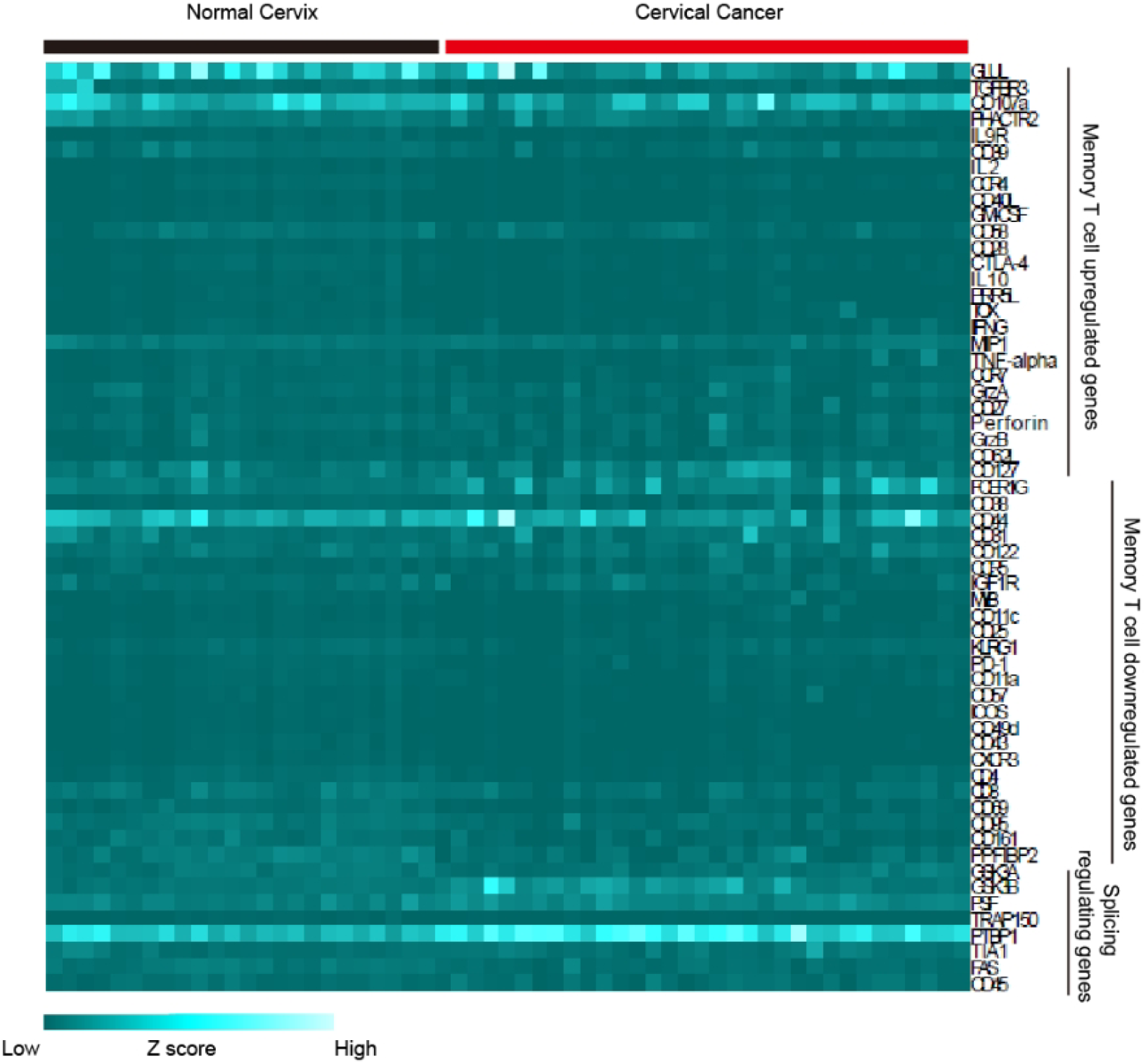
The marker genes related to memory T cells in GSE9750 dataset and the key genes regulating the memory cell marker CD45 alternative splicing signaling pathway were screened and analyzed by heat map.

### 2.3 Exosomal miR-124 regulates T cell immune function by regulating the GSK3-PSF-CD45 alternative splicing pathway

By bioinformatics prediction of the targeting relationship between miR-124 and genes related to immune function, we screened several genes that may be regulated by miR-124 (Table 1). Q-PCR analysis of exosomal miR-124 showed that its expression was significantly reduced in HPV persistent infection, while the expression of its predicted target gene GSK3 in PBMC showed an opposite trend (FIG. 10A, B). miR-124 may act on PBMC by exosomes and cause changes in the expression of the target gene GSK3. The protein expression of GSK3 was found to be negatively correlated with miR-124 (FIG. 10C). GSK3-PSF-CD45 is a member of the regulatory protein of the memory cell differentiation-related marker CD45 alternative splicing, and its molecular pathway GSK3-PSF-CD45 can regulate T cell activation and memory cell differentiation (Figure 5). The mRNA expression of the alternative splicing pathway PSF and TRAP150 was detected in peripheral blood PBMC of HPV persistent infection and control. No significant difference was found (FIG. 11). In summary, miR-124 expression was decreased in blood exosomes of HPV persistent infection and affected the immune response through the CD45 alternative splicing pathway by regulating the target gene GSK3.

**Table 1.** Online prediction results of miR-124 and 3’UTR complementary miRWalk of immune function related genes.

| mirnaid | genesymbol | start | end | bindingp | energy | seed |
| --- | --- | --- | --- | --- | --- | --- |
| hsa-miR-124-3p | IL31 | 729 | 749 | 1 | -18.4 | 1 |
| hsa-miR-124-3p | IL2RA | 2203 | 2220 | 1 | -20.5 | 0 |
| hsa-miR-124-3p | IL11 | 718 | 738 | 1 | -20.5 | 1 |
| hsa-miR-124-3p | GSK3B | 5787 | 5800 | 1 | -20.1 | 0 |
| hsa-miR-124-3p | IL2RB | 2209 | 2229 | 1 | -20.8 | 1 |
| hsa-miR-124-3p | FAS | 3396 | 3414 | 1 | -20.1 | 0 |
| hsa-miR-124-3p | CXCR4 | 1350 | 1367 | 1 | -18.2 | 0 |
| hsa-miR-124-3p | PDCD1 | 1942 | 1954 | 1 | -19.3 | 0 |

**Figure 10.**
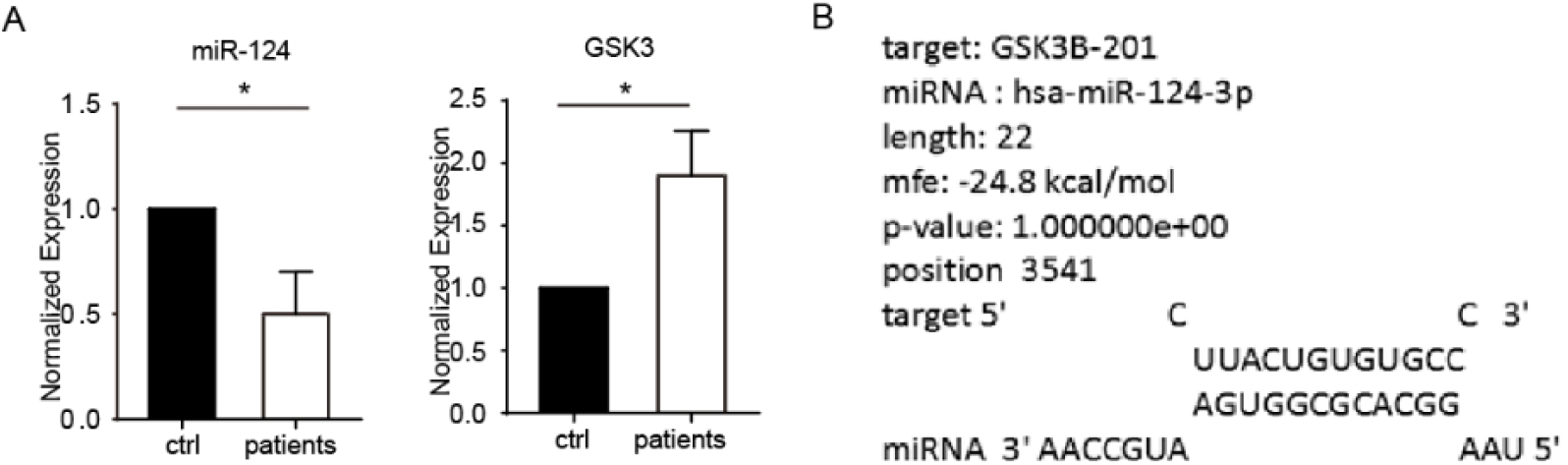
Expression levels of miR-124 and GSK3 in exosomes from peripheral blood of cervical cancer patients and controls without infection detected by Stem-loop Q-PCR (A). And miR-124 and GSK3 bioinformatics target sequence prediction (B).

**Figure 11.**
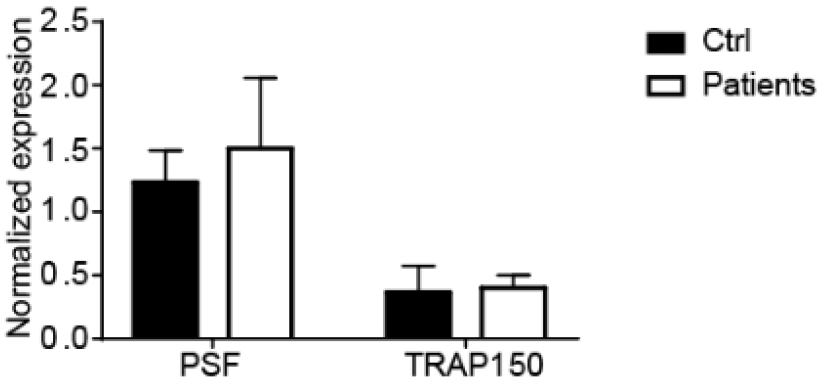
mRNA expression of key genes regulating the memory cell marker CD45 alternative splicing signaling pathway by Q-PCR in peripheral blood PBMC cells from high-risk HPV infected and controls.

## 3 Discussion

In this study, we observed that plasma exosomes derived from persistent HPV infection can diminish the proportion of Th immune memory cells and significantly decrease the expression of miR-124 in exosomes. This suggests that miR-124 plays a regulatory role in immune response and influences the immune clearance of HPV infection. The correlation between exosomal miR-124 expression and persistent HPV infection was confirmed by assessing the levels of exosomal miR-124 in both case and control populations. Furthermore, we validated the impact of exosomal miR-124 on Th immune function through in vitro treatment of PBMC with exosomal miR-124 as well as through overexpression or silencing experiments involving Mir-124. Finally, we elucidated the role of exosomal miR-124 in HPV virus immunity.

Persistent high-risk HPV infection is a crucial risk factor for cervical cancer development. Previous studies have reported an association between genetic polymorphisms in miR-124 and cervical cancer occurrence; however, research investigating the correlation between persistent HPV infection and cervical cancer is lacking. In our investigation, we identified decreased expression of miR-124 within plasma exosomes from individuals with persistent HPV infection. Additionally, we identified three SNPs (rs531564, rs73662598, and rs73246163) that may influence transcriptional regulation and processing mechanisms associated with miR-124 through a case-control study aimed at clarifying their relationship with high-risk HPV persistence. These findings hold significant implications for identifying susceptible populations to high-risk HPV persistence as well as for developing therapeutic vaccines.

The regulatory effect of exosomal miR-124 on the alternative splicing pathway of CD45, a memory cell marker, was demonstrated through in vitro molecular biology experiments. Pathway blocking experiments further confirmed the specific mechanism by which miR-124 regulates Th cell immune regulation, providing crucial targets for future research on HPV vaccine and immunotherapy. However, several challenges remain to be addressed, including the clinical feasibility of applying exosomal miRNA-124 to regulate CD45 alternative splicing in cervical cancer immune regulation. Further investigation into the mechanism and function of exosomal miRNA-124 in regulating CD45 alternative splicing will offer a novel breakthrough and therapeutic strategy for cervical cancer immunotherapy.

## 4 Conclusions

Exosomal miRNA-124 plays a crucial role in the immune regulation of cervical cancer and influences the tumor immune microenvironment by modulating CD45 alternative splicing. We observed a significant downregulation of miRNA-124 expression in cervical cancer tissues, with notable differences compared to normal cervical tissues. Through experimental validation, we discovered that exosomal miRNA-124 can transfer information between cervical cancer cells and immune cells via trafficking within tumor cell exosomes. Further investigations revealed that miRNA-124 regulates the activity and function of immune cells by controlling CD45 alternative splicing. CD45 is a pivotal cell surface protein involved in signaling regulation and activation of T cells, B cells, and natural killer cells. Our findings demonstrate that miRNA-124 interacts with the alternative splicing site of CD45 to regulate its exon splicing, thereby impacting its expression form and function. Additionally, we found that miRNA-124 significantly affects the differentiation and function of immune cells, exerting an important influence on the cervical cancer tumor microenvironment. Specifically, it inhibits anti-tumor capabilities by promoting differentiation into tumor-associated M2 macrophages while also suppressing proliferation and activation of T cells through modulation of immune cell functions, ultimately reducing anti-tumor immune responses within the tumor microenvironment. In summary, exosomal miRNA-124 participates in the immune regulation of cervical cancer through modulation of CD45 alternative splicing. This discovery provides novel insights into mechanisms underlying immune evasion in cervical cancer and establishes a theoretical foundation for future development of therapeutic strategies targeting both miRNA-124 and CD45. However, further studies are required to elucidate specific mechanisms involving miRNA-124 in the occurrence and progression of cervical cancer, as well as its potential and limitations in clinical application.

## Acknowledgments

This study is funded by the Guangxi Science and Technology Department (Gui Ke AD22035164, Gui Ke AD22035149), the Guangxi Education Department (2022KY0494, 2022KY0503), and the National Natural Science Foundation of China (82260423)

